# 41% of Gram-negative bacteria encode a Type Six Secretion System

**DOI:** 10.64898/2026.11.07.764895

**Authors:** Christopher Zhang, Hayley Hassler, William C Ratcliff, Lauren Speare, Brian K Hammer

## Abstract

The type six secretion system (T6SS) is a highly studied, versatile bacterial nanoweapon that is important in bacterial pathogenesis and evolution. Despite extensive study, its prevalence remains poorly quantified. Analyzing 35,467 representative genomes, we find that approximately 41% of genomes in the average well-sampled Gram-negative genus (more than 30 sampled genomes) encode a T6SS. Further, we show that the distribution of the T6SS is patchy at all taxonomic levels. The uneven distribution we found is consistent with context-dependent selection, repeated evolutionary loss, or greater evolutionary lability. Our dataset provides a foundation for studying the drivers of T6SS distribution, and our pipeline enables similar surveys of other secretion systems, gene clusters, and multi-protein complexes across bacteria.

## Introduction

The type six secretion system (T6SS) was initially described in 2006 in *Vibrio cholerae* as a novel protein secretory system with anti-eukaryotic activity (1, 2). Over the following two decades, this versatile nanoweapon has been a central focus of research on pathogen virulence (3–5), interbacterial competition (5–7), and toxin-immunity systems (8–10). The original conception of T6SS as merely an anti-eukaryotic weapon has expanded to antagonism against organisms from every domain of life (3, 11–13), nutrient scavenging (14, 15), host interaction (6, 16, 17), kin discrimination (18, 19), human pathogenesis (20–23), plant symbiosis (24), and even natural transformation (6).

Given the significant influence of T6SS activity on microbial community dynamics, understanding its prevalence across the bacterial domain warrants systematic investigation. To this end, one of the early papers on the T6SS surveyed approximately 400 bacterial genomes and reported that T6SS was present in 25% of Gram-negative bacteria (25). This ∼ 25% figure has been widely cited, references to this statistic appear in approximately one third of the roughly 700 T6SS studies cataloged in PubMed with more than five citations (Fig S1). However, this figure provides a limited view of the distribution of the T6SS, as genomic datasets at the time were heavily skewed toward well-studied pathogenic bacteria.

Recent advances in sequencing technologies and bioinformatic tools have enabled more comprehensive and accurate assessments of T6SS distribution across more diverse microbial populations. Techniques such as the HMm Based UndeRstanding of gene clustERs (HAMBURGER) tool (26, 27) have been used to quantify the abundance of T6SS within well-sequenced strains. New protein imaging technologies have allowed us to more accurately discern important and necessary structures within the T6SS complex (28–30). Increased availability of inexpensive microbial sequencing has greatly expanded the number of samples available for analysis, particularly from environmental sources where the role of T6SS in shaping bacterial populations is less clear (31, 32). In this study, we used the Markov Cluster (MCL) algorithm, a graph-clustering method, to systematically identify T6SSs across over 35,000 representative genomes (>70-fold increase in genome quantity compared to the original study) within the Genome Taxonomy Data Base (GTDB), generating the largest survey of the phylogenetic distribution of the T6SS to date. We use this framework to determine the prevalence of T6SS across Gram-negative bacteria and resolve how T6SS distribution varies across taxonomic scales. Our analyses reveal that T6SSs are remarkably widespread yet heterogeneously distributed across bacterial phylogeny, providing new insight into the ecological and evolutionary forces shaping this competitive mechanism. More broadly, our approach provides a framework for investigating the phylogenetic distribution of other complex, multi-protein bacterial systems.

## Main

### Detection criterion

A major challenge in estimating T6SS prevalence is the lack of a standardized definition of a complete T6SS. In nearly every study, authors use different ad hoc criteria as the operating definition for the T6SS (14, 25, 33– 42). Although many studies agree that *tssA-tssM* and PAAR represent the core T6SS genes, consensus breaks down regarding the necessity of components relevant for accessory function such as genes coding for effectors or regulatory genes. For clarity, we show a complete composite structural model of all of the consensus core genes in Fig. 1.

**Fig 1.**
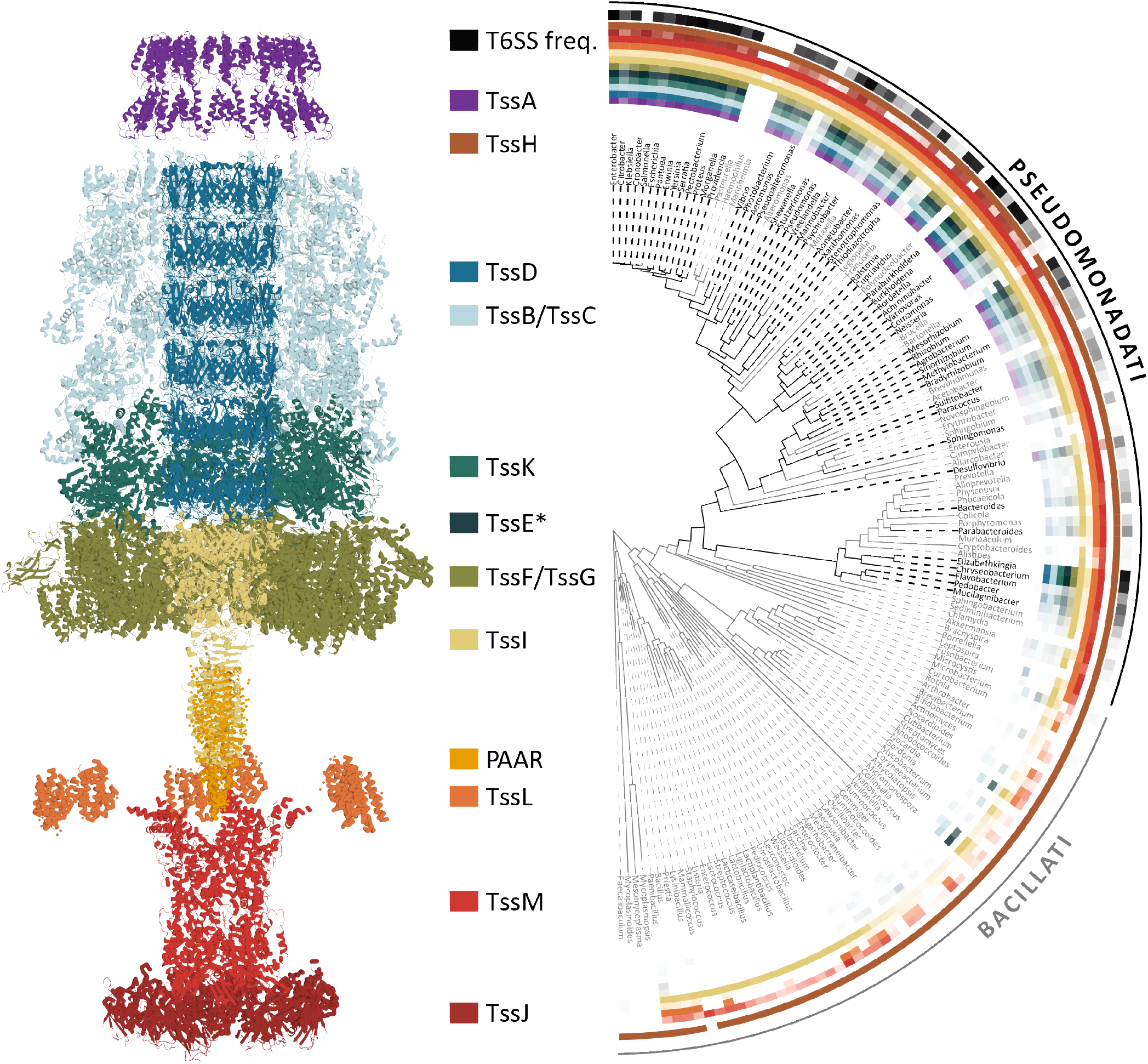
Composite structural model of the T6SS apparatus and phylogenetic distribution of T6SS components across bacterial genera. The structural model was assembled in UCSF ChimeraX by rigid-body fitting of PDB structures into two cryo-EM maps: the 8 Å *Vibrio cholerae* T6SS baseplate/needle reconstruction (EMD-3879) and the *V. cholerae* membrane complex (EMD-0264). Components fitted were TssH/ClpV (3ZRI), TssI/VgrG (4MTK), TssL/TagL (4V3I), TssM and TssJ (6HS7), PAAR (4JIV), TssB/TssC and TssD/Hcp (5OJQ), TssA (6RJU), and TssK/TssF/TssG (6N38). Select subunits of TssB/TssC, TssK, TssF/TssG, and TssL were omitted to provide a cross-sectional view of the interior. A maximum-likelihood phylogenetic tree was constructed from concatenated *rpo*B, *rpo*C, and *rps*J gene sequences and rooted using midpoint rooting, collapsed to genus level by pruning redundant sister taxa sharing identical genus designations. Branch and tip label colors indicate whether the genus has a T6SS prevalence above or below a 20% threshold. Heatmap rings flanking the tree display the percentage of genomes within each genus encoding each core T6SS component, with color intensity proportional to prevalence (white = 0%, full color = 100%). Gene families shown include the inner tube (TssD/Hcp), contractile sheath (TssB/TssC), baseplate subcomplex (TssK, TssF/TssG), spike complex (TssI/VgrG, PAAR), membrane complex (TssJ, TssL, TssM), AAA+ ATPase (TssH/ClpV), and cap protein (TssA). TssE, the gp25-like baseplate wedge protein, was not modeled because it is absent from the TssK/TssF/TssG cryo-EM structure (6N38) and its position within the baseplate remains uncertain (28). The outermost ring displays the raw T6SS prevalence per genus.

To address the ambiguity in T6SS composition, we selected five criteria spanning the range of definitions in common use, chosen for their recent usage, their importance to the field of T6SS, and their varying levels of stringency. We named these criteria after the first author of the article that introduced them:

1. The Bingle criterion: the original paper from which the 25% figure was calculated requires both *tssB* and *tssC* (25).
2. The Boyer criterion: one of the first genome-wide surveys of the T6SS required at least five distinct T6SS genes in a single cluster, with no more than four intervening genes between consecutive hits (38).
3. The Suria criterion: a recent prevalence survey utilized a custom application of the HAMBURGER tool to identify T6SSs among squid symbionts. In their analysis, they required at least four distinct core structural genes (*tssA–tssM*) clustered within ten intervening non-T6SS genes (42).
4. The Stubbusch criterion: a large recent analysis of environmental marine taxa had a stringent definition requiring more than two-thirds of the fourteen core components (*tssA– tssM* and PAAR, the homolog of EvpJ) for a genome to be considered T6SS positive (14).
5. The TXSScan criterion: the widely used TXSScan models for MacSyFinder require at least eleven of the fourteen core genes (*tssA-tssM* and *evpJ*) in a single cluster, with no more than twenty intervening genes between hits; we applied these thresholds to our gene calls rather than running MacSyFinder itself (43).

### Well-sampled Gram-negative genera have an average of 41% T6SS prevalence

We took a stratified sample of bacterial genomes from the GTDB r226 release as our representative dataset of bacteria. The GTDB is a large, robustly curated set of bacterial genomes drawn from RefSeq and Genbank (32). Each GTDB release includes a mixture of well sequenced bacterial genomes and draft genomes from uncultured cells, such as those obtained through metagenomics, allowing for our analysis to reflect natural biodiversity more accurately. Our stratification approach reflects this dichotomy: for well sampled genera (greater than 500 total samples in the database) we sampled 100 random genomes, prioritizing higher quality “genome representatives.” For poorly sampled genera (all other genera), we took 10% of the genomes in each genus, again prioritizing higher quality “genome representatives.” We removed any genomes that had not already undergone gene-finding. This sampling strategy yielded 35,476 bacterial genomes that spanned the entire bacterial phylogeny. Our method is enumerated in the methods section and explicitly shown in the supplementary code.

Concurrently, we searched the predicted proteomes of all sampled genomes with the T6SS protein sequences in the Se-cReT6 database (44) using DIAMOND, and grouped the hits into homologous gene clusters with the Markov Cluster algorithm (MCL). Each cluster was annotated by the SecReT6 sequences it contained, which identified the T6SS proteins present in each genome. From these data, we created an expansive lookup table of all detected T6SS proteins, along with their genomic locations for the genomes that could meet a position-based criterion (provided in our online code/data repository).

We cross-referenced our lookup-table with our T6SS criteria to generate T6SS+/-results for all 35,467 genomes under each criterion, and compared every pair of criteria against each other. The Boyer, Stubbusch, and Suria criteria agreed closely with one another (percent agreement 96–98%, Co-hen’s *κ* = 0.84–0.93), whereas agreement was lower for the Bingle criterion, which requires only two genes(*κ* = 0.72– 0.83), and for the strict TXSScan thresholds (*κ* = 0.55–0.80) (Fig. S2).

Next, we assessed how well each criterion recovers known systems while avoiding false positives, using a blind validation set of 187 genomes: 104 with experimentally validated T6SS containing genomes from the Secret6 database (44) and 83 randomly chosen negatives also provided by Secret6. Against this dataset, the Suria and Boyer criteria performed the best (Sensitivity = 100% and 98%, Specificity = 88% and 89%, MCC = 0.90 and 0.88) whereas the Stubbusch criterion was slightly more specific but less sensitive (Sensitivity = 79%, Specificity = 92%, MCC = 0.70). The Bingle and TXSScan criteria fared worse, with the Bingle criterion over-calling T6SSs (63% specificity) and the TxSScan being overly stringent (34% sensitivity) (Fig. S3).

Because each of these criteria captured different notions for what a complete T6SS system is, and because each criteria had different strengths and weaknesses, we defined a consensus criteria that incorporated the Boyer, Suria, and Stubbusch criterion together as they were the best performing. The combined criteria would nominate an genome as T6SS+ when at least two of the constituent criteria agreed. This consensus call recovered 98% of validated systems with 89% specificity (MCC = 0.88); we used it as a baseline metric for subsequent analyses.

To account for uneven sampling across genera, we first calculated T6SS prevalence within each genus as the proportion of its sampled genomes encoding a T6SS, then averaged these values across genera so that each genus contributed equally. An enumerated list of all genera across all criteria is provided in the online data repository. Then, we cross-referenced our dataset with BacDive’s dataset on Gram-status. Weighting each genus equally, we find that T6SS prevalence averages 41% across well-sampled Gram-negative genera (more than 30 sampled genomes, n = 77). This finding was robust to the choice of criterion (40-42% across Suria, Stubbusch, and Boyer criteria individually; Fig. S4) and to how GTDB genera were matched to Gram-stain records (41% when polyphyletic GTDB genera such as *Pseudomonas_E* were assigned to the Gram stain of their parent genus). It is important to note that we also observe a strong frequency-dependent effect in our prevalence calculation; higher sampled bacteria have a higher prevalence of T6SS (Fig. S4). Even at its lowest, the consensus criterion finds that the prevalence of the T6SS is over 29%.

This genus-averaged prevalence exceeds the widely cited 25% estimate (25) and suggests that T6SSs may play a more central role in bacterial ecology than the existing literature implies. This 41% approximation describes a genus-level prevalence and not a species or strain specific one and only represents well-sampled bacteria (over 30 sampled genomes). Our result should be interpreted as a measure of the phylogenetic distribution of the T6SS at a genera level rather than its frequency among strains or its in absolute quantity the environment.

### T6SS is patchy at all taxonomic levels

In the previous section, we showed that T6SS prevalence averages 41% across well-sampled Gram-negative genera. This average describes the overall prevalence of the T6SS but not how it is distributed among related lineages. This question is important as the presence or absence of the T6SS within individual strains have been very difficult to predict, even among well studied strains of bacteria (42, 46–50). Anecdotally, simply knowing the lineage of a bacterium does not predict T6SS utilization or function; many studies find high variance in T6SS down to the strain level (48, 51–57). With our dataset, we can interrogate this question of T6SS distribution at a finer grain to quantify the prevalence of the T6SS among closely related strains.

At its broadest level, our findings show that with few exceptions, the T6SS is restricted to Gram-negative bacteria (Fig. S5). Across the 1,222 Gram-negative genera with a Gram stain recorded in BacDive, the mean genus T6SS prevalence was 28.6%, compared with 0.2% across 572 Gram-positive genera (each genus weighted equally; Fig. S5A). The gap widens when only well-sampled genera (more than 30 sampled genomes) are considered: mean prevalence rose to 41.5% for Gram-negative genera (*n* = 77) but stayed at 0.1% for Gram-positive genera (*n* = 31), and 81% of well-sampled Gram-negative genera contained at least one T6SS-positive genome, compared with 6% (2 of 31) of Gram-positive genera (Fig. S5B). This finding is consistent with its dual membrane-anchored architecture (46) and has been commonly assumed within the T6SS literature (7, 58, 59). However, within Gram-negative bacteria, as many others have found, the presence or absence of T6SS is much more difficult to predict. To quantify the level of within-lineage diversity, we selected a metric called the Gini-coefficient.

We noticed that our dataset was similar to some datasets more commonly found in economics, specifically economic inequality datasets. Individuals in this dataset (analogous to genera in our dataset) all have a proportion of the total wealth (percent T6SS+), some individuals have more (high T6SS+ proportion) and some individuals have less (low T6SS+ proportion). In these datasets, the Gini-coefficient is used to quantify the amount of inequality by measuring the ratio of the area between the Lorenz curve and the line of equality to the total area underneath the line of equality (or half of the relative mean absolute difference) (Witte-bolle *et al*. contains a concise explanation of the calculation of the Gini-coefficient in the biological sciences (60)). The Gini-coefficient measures inequality among values of any frequency distribution and has been previously used to calculate ecological community evenness (60). Helpfully, this statistic provides a singular metric to characterize patchiness, making it a useful comparative tool in our dataset.

In our dataset, a Gini-coefficient of 0 indicates that all of the genera within a higher taxonomic rank have the same T6SS prevalence. For example, if every Enterobacteriaceae genus were 60% T6SS+, the family would have a Gini coefficient of 0. If the Enterobacteriaceae family had maximal unevenness, e.g. only one genus within Enterobacteriaceae contained the T6SS and all other genera do not contain any T6SS, this family would have a Gini-coefficient approaching 1.

Because genus-level prevalences are between 0 and 1, the Gini-coefficient of a family with mean T6SS prevalence *µ* is bounded above by *G* ≤ 1 − *µ*, with equality if and only if every genus is either entirely T6SS+ or entirely T6SS-. Therefore, the distance between a family’s Gini-coefficient and the upper bound described by the line *y* = 1 − *x* measures how close the family comes to this maximally bimodal state, with the distance from the line describing the maximum number of intermediate values. We calculated the Gini-coefficients for each bacterial family with at least five genera and at least one T6SS+ genus (Fig. 2A).

**Fig 2.**
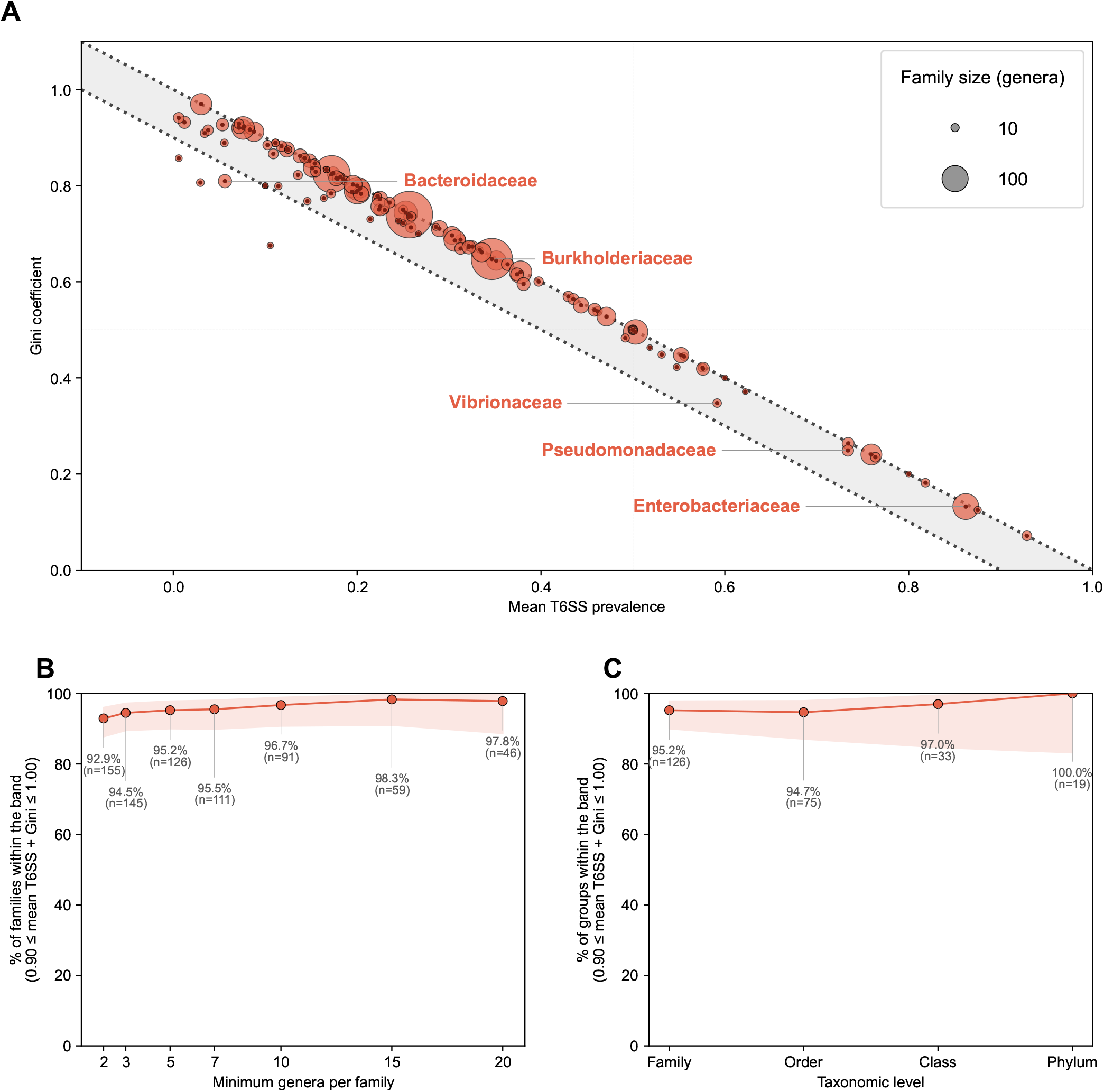
T6SS is patchy in Gram-negative bacteria. (**A**) Gini coefficient of T6SS prevalence versus mean T6SS prevalence for each Gram-negative bacterial family containing at least five genera and at least one T6SS-positive genus (*n* = 126 families). For each family, T6SS prevalence was calculated per genus as the proportion of its sampled genomes called T6SS-positive by at least two of the Suria, Stubbusch, and Boyer criteria, and the Gini coefficient was computed across all constituent genera, including T6SS-absent genera (prevalence = 0); a coefficient of 0 indicates uniform prevalence across genera and 1 indicates concentration within a single genus. Point area is proportional to the number of genera in the family; the black dot marks each family’s center. Family Gram status was assigned as Gram-negative when the majority of Gram-classified genera were Gram-negative, using genus-level Gram assignments from BacDive (45). Dotted lines mark *y* = 1 − *x* and *y* = 0.9 − *x*; the shaded band between them contains families whose mean prevalence and Gini coefficient sum to between 0.90 and 1.00, corresponding to families composed almost exclusively of uniformly T6SS-positive and uniformly T6SS-absent genera, with few genera at intermediate prevalence. (**B**) Percentage of Gram-negative families falling within the band in (A) as a function of the minimum genus-count threshold. The denominator at each threshold is all Gram-negative families with at least that many genera and at least one T6SS-positive genus, and a mean prevalence below 1; error bars/labels report the count *n* at each threshold. The fraction is broadly stable across minimum-genera thresholds. (**C**) Percentage of Gram-negative groups within the band, computed at each taxonomic rank from family upward (family, order, class, phylum). At every rank, the Gini coefficient is measured across genera (each genus weighted equally, absent genera included), so the unit of measurement is held constant as the aggregating rank increases. Shaded bands in (B) and (C) denote 95% Wilson score confidence intervals; *n* per point is annotated. The patchiness of the T6SS is broadly stable across taxonomic rank.

We find that the vast majority of families cluster near this upper bound with 95.2% of families (120 of 126) having a Gini-coefficient within 0.1 of the maximum possible value given its mean T6SS prevalence. Fig. 2A reveals that within families, and at every level of mean T6SS prevalence, there are many genera in which the T6SS is either highly prevalent or entirely absent, but very few genera with intermediate prevalence.

Additional sensitivity testing shows that this trend is broadly stable across minimum genus-count thresholds from 2-20 (92.9–98.3%; Fig. 2B) and across taxonomic level (94.7– 100%; Fig. 2C) suggesting that our result is not an artifact of the sampling threshold, though we note that genera excluded by stricter thresholds are disproportionately understudied lineages whose true patchiness remains uncertain.

This patchiness raises a question that the prevalence estimate alone cannot answer. Previous studies in some T6SS lineages have found that T6SS imposes negligible metabolic cost under a variety of expression conditions (61, 62), so widespread loss across certain lineages is not easily explained by energetic burden. One possible explanation for this finding is that the cost-benefit balance of maintaining an interbacterial weapon depends heavily on community and resource contexts: population density, the identity of neighboring taxa, and the local diversity of competing effector arsenals contribute to the presence or absence of the T6SS (6, 8, 63).

## Conclusions

Across 35,467 genomes, we estimate that approximately 41% of genomes in well-sampled Gramnegative genus encode a T6SS, well above the 25% figure that has anchored the field since 2008 (25). The T6SS is also strongly patchy at every taxonomic rank; genera tend either to carry the system at high prevalence or to lack it entirely, with few genera in between. Together, these results suggest that the T6SS is both more common and more evolutionarily labile than previously appreciated.

Previous surveys of T6SS prevalence focused on specific bacterial groups (35, 39, 41, 42, 64, 65), analyzed smaller datasets (33), measured environmental rather than phylogenetic distribution (14, 34), or did not specify which genes constitute a T6SS (24). To date, our analysis comprises the highest quantity of samples, examined across the widest taxonomic tree, and analyzed with the most up-to-date methods yet. We also provide two resources for the community: a public dataset of T6SS gene calls and genomic locations for 35,391 genomes, which will support more refined estimates as genome databases grow, and a detection pipeline that generalizes to other multi-protein systems.

Several challenges still complicate the assessment of T6SS prevalence. First, the term “Gram-negative” is an operational category based on staining properties rather than a mono-phyletic clade, and many bacteria do not fit the Gram-binary classification system. Because we assigned Gram status from BacDive records, genomes from uncultured lineages lack a Gram assignment; we include them in taxonomy-wide analyses but exclude them from Gram-negative estimates. Second, no detection criterion is ideal: stringent criteria may miss divergent or minimal systems, while permissive criteria may count incomplete or nonfunctional clusters. Criteria that count genes anywhere in the genome (Stubbusch) can assemble an apparent system from scattered homologs; indeed, 23% of Gram-positive genomes, which canonically lack a T6SS, carry four or more core-gene homologs (Fig. S7E). Conversely, criteria that require genes to be clustered (Suria and Boyer) may miss systems whose loci are split across the genome or across contigs of fragmented assemblies. To mitigate these issues, we required agreement between at least two of the three best-performing criteria and confirmed that each of these criteria alone gives a similar estimate (Fig. S4). Third, T6SS prevalence depends on how well a genus is sampled. Our sampling strategy could not represent highly and sparsely sampled genera evenly, as we analyzed 10% of the genomes in genera with up to 500 genomes and 100 genomes from each larger genus. Mean prevalence across Gram-negative genera rises from 28.6% across all genera to 41.5% among genera with more than 30 sampled genomes (Fig. S4, Fig. S5A). Detection may contribute to this bias: genera with very few sampled genomes are largely uncultured lineages represented by metagenome-assembled genomes (Fig. S6), so the losses from fragmented assemblies may fall disproportionately on poorly sampled genera, in which case their true prevalence is higher than we report. The pattern could also be biological. Many of the most heavily sampled bacteria come from environments of high bacterial density, such as the human microbiome, the rhizosphere and biofilm mats, whereas poorly sampled genera may come disproportionately from environments with fewer bacteria, where a system used in interbacterial competition may be less common. Nevertheless, even when poorly sampled genera receive the greatest weight, the consensus estimate (28.6%; 24.7–30.6% across its three component criteria) is at or above the commonly cited figure of about 25%, and it reaches 47.5% when only genera with more than 40 sampled genomes are considered (Fig. S4).

Our approach can map other multi-protein systems, including other secretion systems, microcompartments, competence machinery, and flagella, as well as gene sets that function in conjunction, such as bacterial immune systems, quorum sensing circuits, secondary-metabolite biosynthetic gene clusters, and sporulation pathways. It would even be possible to tie related bacterial components together evolutionarily, for example, we can begin to disentangle the complex relationship between conjugation machinery and bacterial immune machinery. For the T6SS specifically, our dataset may be large enough to test, for the first time at scale, the proposed evolutionary links among phage tails, extracellular contractile injection systems, pyocins, and the T6SS. By integrating environmental metadata, scientists can begin to identify the ecological factors that impact T6SS prevalence and comparative genomics could reveal genes that co-evolve with the T6SS cassette.

Together, these resources position the community to move beyond the question of how common the T6SS is, and toward the more interesting question of why its distribution looks the way it does.

## Methods

### Literature survey of T6SS prevalence claims

To quantify how often the T6SS literature makes claims about how widespread the system is, we searched PubMed Central (PMC) through NCBI on November 11th, 2025 for articles containing any of the terms “T6SS”, “Type Six Secretion System”, “Type VI Secretion System” or “Type 6 Secretion System”, joined with OR and searched across all fields.

This broad search returned 6,500 articles, including many that mention the T6SS only in passing. The list of PMC identifiers was exported, and the full-text XML of each article was downloaded from PMC with the Entrez E-utilities (Biopython; *efetch*, database *pmc*), with requests rate-limited to NCBI’s guidelines and failed downloads retried. The PubMed identifier (PMID) of each article was read from its XML record (6,493 of 6,500 articles had one).

Citation counts and publication years were obtained from NIH iCite (exported November 13, 2025) and joined to the articles by PMID. Articles cited more than five times (n = 4,142) were retained so that the survey reflects work that has been read and cited. To keep only papers whose subject is the T6SS, the title and abstract of each remaining article were parsed from the XML and searched with case-insensitive regular expressions for “T6SS”, “type VI secretion system”, “type 6 secretion system” or “type six secretion system”. Articles with none of these terms in the title or abstract were excluded, leaving 736 T6SS papers published between 2007 and 2025.

For each paper, the abstract and the introduction were extracted from the XML. The introduction was taken as the section tagged as an introduction, or titled “Introduction” or “Background”; when no such section was tagged, the first body section was used if its title contained “intro” or “background”. Methods, results and discussion sections were not searched, so that only statements framing the field were counted. The extracted text was split into sentences, and each sentence was tested against a set of case-insensitive regular expressions describing prevalence claims about Gram-negative bacteria or Proteobacteria. These covered (i) the quantitative estimate (e.g. “25% of Gram-negative bacteria”, “a quarter of Gram-negative bacteria”); (ii) distribution claims (e.g. “widespread among”, “widely distributed in”, “found in many”, “broad distribution in”); (iii) frequency claims (e.g. “common in”, “prevalent among”, “ubiquitous in”, “abundant in”, “the majority of”, “most Gram-negative bacteria”); and (iv) conservation claims (e.g. “conserved among”). A final pattern matched any sentence that contained both a prevalence term and “Gram-negative” or “Proteobacteria”. A paper was classified as making a prevalence claim if at least one sentence matched; matched sentences were saved for inspection.

Papers were summarized by publication year (Fig. S1A) and by citation count in four bins: 6–20, 21–50, 51–100 and more than 100 citations (Fig. S1B). For each bin, and for all papers combined, the percentage of papers making a claim is reported with a 95% Wilson score interval (statsmodels).

### Sampling, Labeling, and Analysis Strategies

Genomes were sampled from the Genome Taxonomy Database (GTDB) r226 release. Because the GTDB is dominated by a small number of heavily sequenced genera, we used a stratified design that caps sampling effort in large genera while retaining coverage of sparsely sampled, largely uncultured lineages. Genera were classified as large (≤500 genomes) or small (500 genomes); this threshold separates well-studied model genera from lineages known mainly from metagenome-assembled genomes (Fig. S6). From each large genus we sampled 100 genomes, and from each small genus we sampled 10% of its genomes (rounded down, minimum of one). We prioritized ‘GTDB representative genomes’ for each genera, these are high quality genomes that are meant to represent the centroid of a species cluster. When a genus contained enough representatives, the sample was drawn entirely from the representative pool; otherwise, all representatives were taken and the remainder was filled by random draws from non-representative genomes in the same genus.

This sampling strategy gave a total of 58,609 genomes. Of these sampled genomes, 35,476 (from 17,257 genera) had an NCBI protein annotation and were analysed; the remaining 23,133 had no annotated proteins available. All genomes were assigned to genus and higher ranks using GTDB r226 taxonomy after stripping RefSeq/GenBank prefixes (RS_, GB_) from accessions. GTDB genus names, including polyphyly suffixes (e.g., *Pseudomonas_E*), were retained as distinct genera.

### Genome database curation and homologous gene clustering

Protein sequences for the sampled genomes were downloaded from NCBI RefSeq and GenBank (accessed July 22, 2025). All predicted proteomes were concatenated into a single reference database built with DIAMOND v2.1.8.162 (66). The database was queried with DIAMOND blastp using all T6SS protein sequences in SecReT6 (44) (E-value ≤ 10^−5^, -sensitive mode, to recover divergent homologs). The SecReT6 query sequences were also included in the reference database as positive controls for annotation.

Hits were grouped into homologous gene clusters with the Markov Cluster algorithm (MCL; scheme 2, inflation 1.8) (67), parameters previously shown to be robust against false positives (68). Each cluster was annotated by the SecReT6 control sequences it contained. This produced per-genome calls for 24 T6SS protein families: TssA–TssP, VgrG, PAAR, ClpV, Hcp, TagL, IglA/TssB, and fused TssB/TssC and TssF/TssG annotations. Calls for each protein were recorded together with their accession, giving a lookup table of 603,716 protein calls across 35,391 genomes that carried at least one hit.

### T6SS detection criteria

Each genome was classified as T6SS-positive or T6SS-negative under each criterion. Annotations were first mapped to eighteen canonical genes: synonyms were merged (clpV to tssH, vgrG to tssI, hcp to tssD, iglA/tssB to tssB). Fused tssB/tssC or tssF/tssG calls were counted as whichever protein was missing. Core components were defined as tssA-tssM and PAAR, tssN-tssP and tagL were considered T6SS proteins but not core components. Each gene was counted once per genome regardless of copy number. For position-based criteria, every coding sequence on a contig was numbered by start coordinate from the genome’s NCBI GFF3 annotation, and distances were measured in quantity of genes between. GFF annotations were retrieved only for genomes that could satisfy a position-based criterion (at least four core or five canonical genes; 19,914 genomes). 9 total genomes had no annotation available and were treated as missing for those criteria.

- **Bingle**: TssB and TssC both present; IglA/TssB counted as TssB, and a fused TssB/TssC call satisfied both (25).
- **Boyer**: at least five distinct canonical genes in a single cluster, with no more than four intervening genes between consecutive hits (38).
- **Suria**: at least four distinct core genes within a single cluster on one contig, with consecutive hits no more than ten genes apart (42).
- **Stubbusch**: at least ten of the fourteen core genes, i.e., more than two-thirds (14).
- **TXSScan**: at least eleven of the fourteen core genes in a single cluster, with no more than twenty intervening genes between consecutive hits (43).
- **Consensus**: T6SS-positive when at least two of the Suria, Stubbusch, and Boyer criteria were positive, T6SS-negative when at least two were negative, and missing otherwise (9 genomes).

### Concordance and validation of detection criteria

To estimate how well each criterion recovers known systems, we built a blind validation set of 187 genomes: 104 genomes carrying experimentally validated T6SSs in SecReT6 (44) (All validated sequences with protein annotations available) as well as 83 genomes drawn from a negative database within SecReT6 (remainder of 100 randomly drawn genomes with protein annotations available). These genomes were annotated with the same clustering pipeline, and each criterion’s calls were compared with the labels to compute sensitivity, specificity, precision, Matthews correlation coefficient, and 95% Wilson score intervals. Genomes with a missing call for a position-based criterion were excluded from that criterion’s denominator. Paired differences between criteria were tested with exact McNemar tests (Holm-corrected). Criterion parameters were fixed on this set before being applied to the full sample.

### Genus-level T6SS prevalence

For each genus and criterion, T6SS prevalence was calculated as the proportion of sampled genomes classified as T6SS-positive, with 95% Wil-son score confidence intervals (statsmodels v0.14.5). For all downstream analyses, each genome was assigned a consensus call where a genome was called T6SS-positive when at least two of the Suria, Stubbusch, and Boyer criteria called it positive, and negative otherwise. Genomes without two agreeing calls (when a position-based call was missing n=9) were excluded from the denominators. Genus prevalence was defined as the proportion of a genus’s sampled genomes with a positive consensus call, with a 95% Wilson score confidence interval.

To test whether the prevalence estimate depends on the criterion, we repeated the genus-level calculation with each criterion alone and compared the mean Gram-negative genus prevalence with the consensus (Fig. S4). To test the dependence on cutoffs, we re-called each count-based criterion at every cutoff from the per-genome scores (core genes in the genome for Stubbusch, and the largest gene cluster for Suria, Boyer and TXSScan-T6SSi), and recorded the mean Gram-negative genus prevalence, the percentage of Gram-positive genera with at least one positive genome (as a false-positive control), and the sensitivity and specificity on the validation set (Fig. S7).

### Gram-stain assignment

Gram-stain records for 15,691 strains were exported from BacDive (45) (advanced search, March 17, 2026). Each strain was assigned to a genus using the first word of its species name. A genus was assigned a Gram status when all of its records agreed, or when one status outnumbered the other by more than 10-fold. Genera with more balanced conflicting records (n = 54) were left unassigned. This produced Gram assignments for 2,334 BacDive genera. These were joined to the GTDB genera by exact genus name, giving 1,794 genera with both a Gram assignment and T6SS data (1,222 Gram-negative and 572 Gram-positive). GTDB polyphyly suffixes were not stripped before this join, so suffixed GTDB genera (e.g., *Pseudomonas_E*) were not matched to BacDive records and were excluded from Gram-specific estimates. As a sensitivity analysis, we repeated the join after removing GTDB polyphyly suffixes, so that each suffixed GTDB genus inherited the Gram stain of its BacDive parent genus while remaining a separate genus in all averages. This added 541 genera (306 Gram-negative, 235 Gram-positive; 2,335 in total) and changed the mean Gram-negative genus prevalence from 41.5% to 41.4% (genera with more than thirty genomes).

### Within-lineage patchiness

Heterogeneity of T6SS prevalence within a lineage was quantified with the Gini coefficient computed across that lineage’s constituent genera:

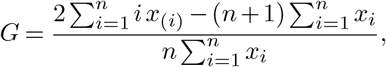

where *x*_(1)_ ≤ … ≤ *x*_(*n*)_ are the ordered genus prevalences and *n* is the number of genera, each genus was weighted equally. Lineages without any T6SS-positive genus were excluded, because *G* is undefined when all *x*_*i*_ = 0.

For prevalences bounded on [0, 1] with mean *µ, G* ≤ 1− *µ*, equality holds only when every genus is either entirely T6SS-positive or entirely T6SS-negative. We therefore classified a lineage as patchy when 0.9 ≤ *µ* + *G* ≤ 1, i.e., when its Gini coefficient fell within 0.1 of the maximum attainable given its mean prevalence.

The primary analysis (Fig. 2A) included Gram-negative families with at least five genera and at least one T6SS-positive genus (*n* = 126). For sensitivity analysis (Fig. 2B), the minimum genus count was varied from 2 to 20. To test dependence on taxonomic rank (Fig. 2C), the analysis was repeated at order, class, and phylum level, again computing *G* across genera so that the unit of measurement was held constant, with a minimum of 5 genera and at least one T6SS-positive genus per group. Confidence intervals on the fraction of patchy lineages are 95% Wilson score intervals.

To assess the presence and distribution of the T6SS across bacteria, we selected 330 genomes representing 165 different bacterial genera of interest. Two genomes per genus were incorporated to both ground the placement of the taxa on the tree and, if present, to highlight potential taxonomic uncertainty within genera. We inferred this phylogeny using a concatenated alignment of the *rpo*B, *rpo*C, and *rps*J (uS10) genes from each genome as these genes have been shown to accurately infer both intra- and inter-genus phylogenetic relationships (69, 70). The concatenated alignment was built using Clustal-Omega v1.2.4 (71). A cluster size of 20 was specified and the alignments were repeated for 3 iterations to ensure maximum accuracy. Using this alignment, we constructed a maximum-likelihood phylogeny of all 330 genomes using IQ-TREE v2.2.2.7 with 1,000 ultrafast bootstrap replicates, specifying the LG+R10 site rate model following optimal model selection by ModelFinder (72–74). Presence and absence of T6SS genes were then mapped to the resulting phylogeny using ggtree and ggplot2 in R version 4.1.2 (75–77).

## Software and code availability

Sampling, classification, phenotype integration, and patchiness analyses were performed in Python 3.9.21 using pandas, polars, NumPy, statsmodels, and matplotlib. Genome annotations (GFF3) were retrieved with the NCBI Datasets command-line tool v18.40.0. All code is available at https://github.com/hhassle/Zhang_et_al_2027.

## ACKNOWLEDGEMENTS

This work was supported by an Department of Education GAANN fellowship P200A210046 to CZ, a NIH NIGMS grant R25GM138030 to WCR, and an NIH NIGMS grant R35GM165598 to LS.

## Supplementary Information

**Fig S1.**
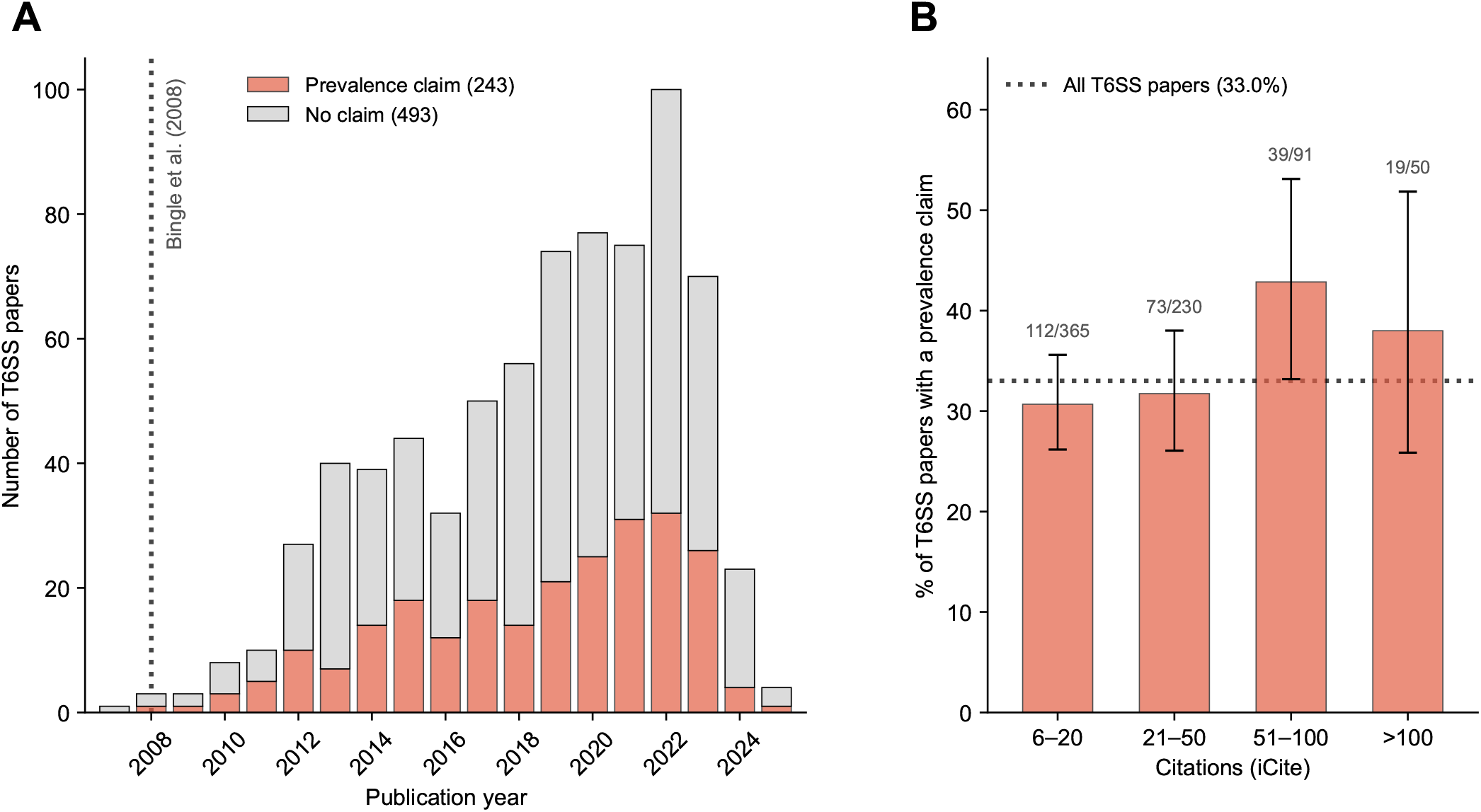
A third of T6SS papers make claims about the widespread nature of the T6SS. Abstracts and introductions of 736 T6SS papers in PubMed Central cited more than five times (iCite) were searched for statements about how widespread the T6SS is. (A) Papers per publication year, split by whether they make a prevalence claim (243) or not (493); the dotted line marks Bingle *et al*. (2008). The decline in 2024 and 2025 reflects the citation filter rather than fewer T6SS studies: of all articles returned by the search, 26.4% from 2024 and 2.9% from 2025 had more than five citations by November 2025, compared with 84–91% of those from 2017 to 2021. (B) Percentage of papers with a prevalence claim by citation count; error bars are 95% Wilson score intervals, numbers above the bars give papers with a claim over papers in the bin, and the dotted line is the rate across all papers (33.0%).

**Fig S2.**
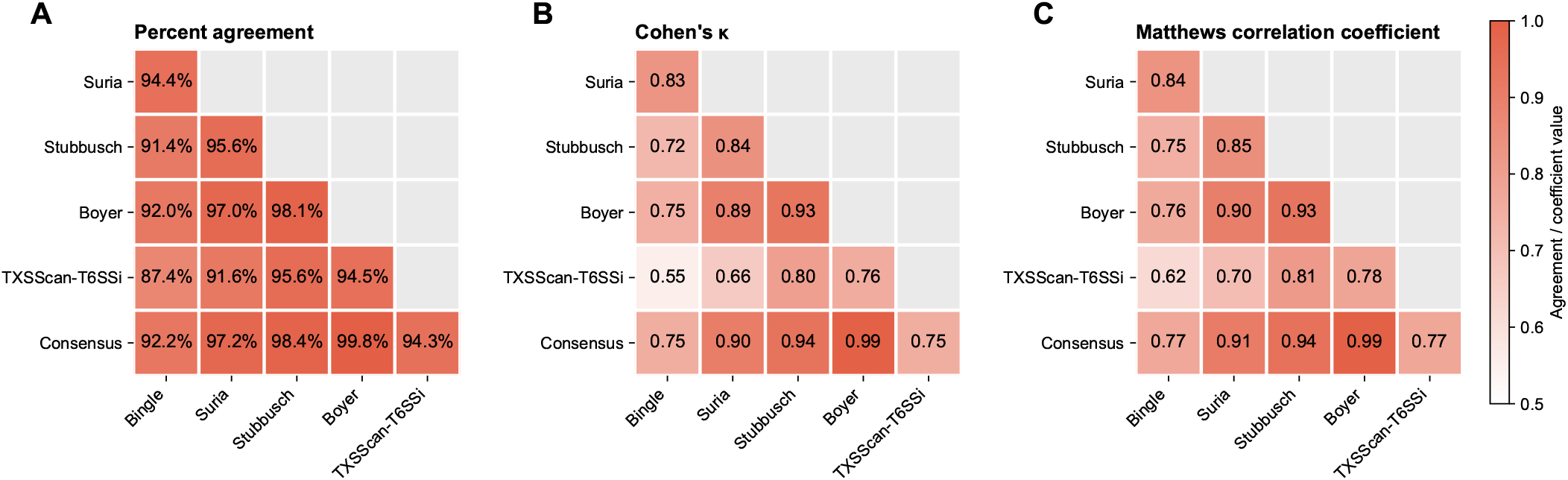
Concordance between T6SS detection criteria. Pairwise comparison of the five detection criteria and the consensus call across 35,467 genomes with a call under every criterion: (A) percent agreement, (B) Cohen’s *κ*, and (C) Matthews correlation coefficient. The color scale runs from 0.5 (white) to 1.

**Fig S3.**
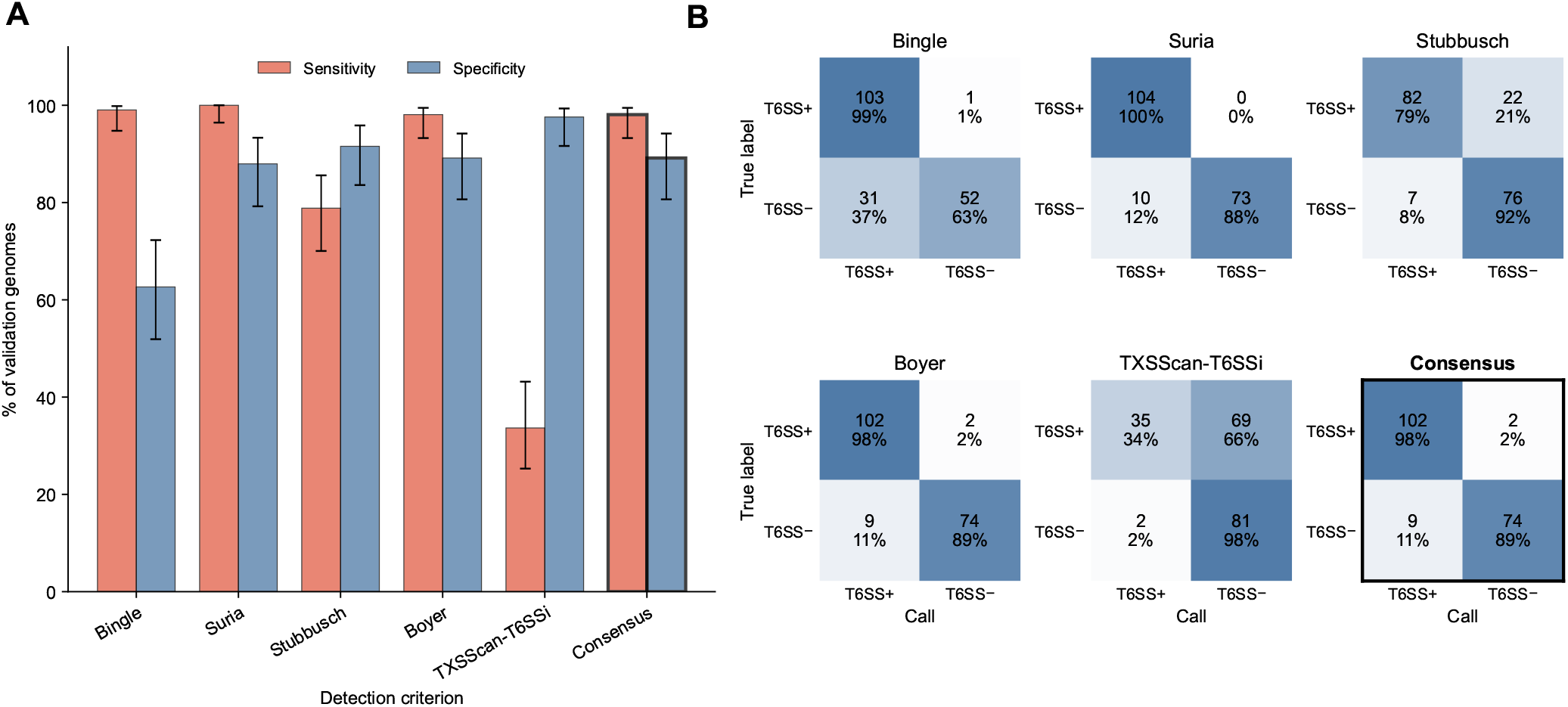
The T6SS detection criteria perform well on a blind validation set. 187 genomes: 104 with experimentally validated T6SSs (SecReT6) and 83 randomly chosen genomes. (A) Sensitivity and specificity of each criterion and the consensus call, with 95% Wilson score intervals; the consensus, used for all downstream analyses, is outlined. (B) Confusion matrices for each criterion and the consensus: rows are the true labels and columns the calls. Each cell gives the number of genomes and its percentage of the row; the top-left cell is the sensitivity and the bottom-right cell the specificity; shading follows the row percentage.

**Fig S4.**
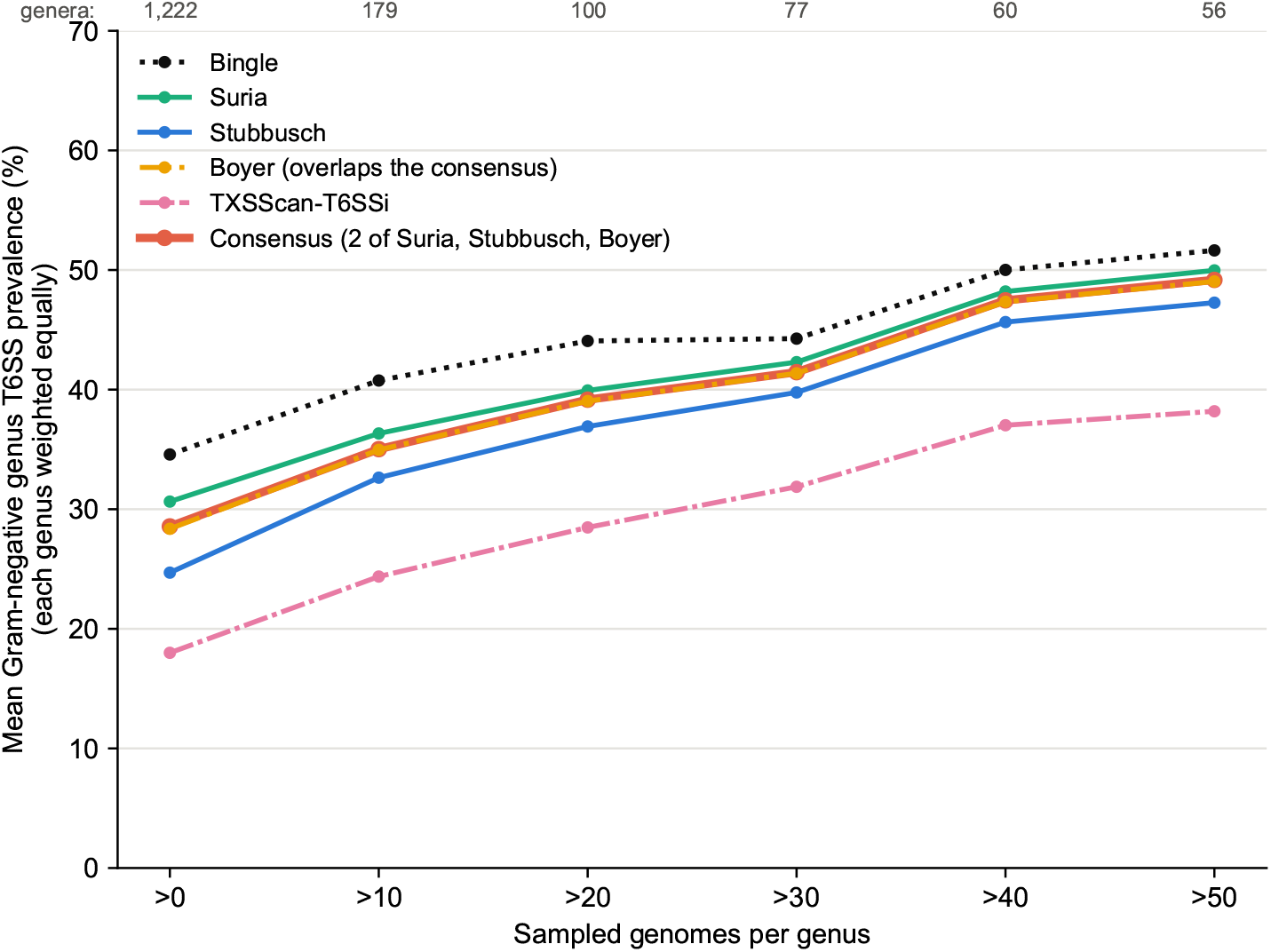
The prevalence estimate does not depend on the detection criterion. Mean T6SS prevalence across Gram-negative genera (each genus weighted equally) under each criterion and the consensus, as genera with few sampled genomes are excluded (cutoffs in steps of ten genomes); the number of genera remaining is given above the plot. Every criterion that combines gene content with gene order gives a similar estimate, and all rise as poorly sampled genera are excluded. The Boyer line overlaps the consensus.

**Fig S5.**
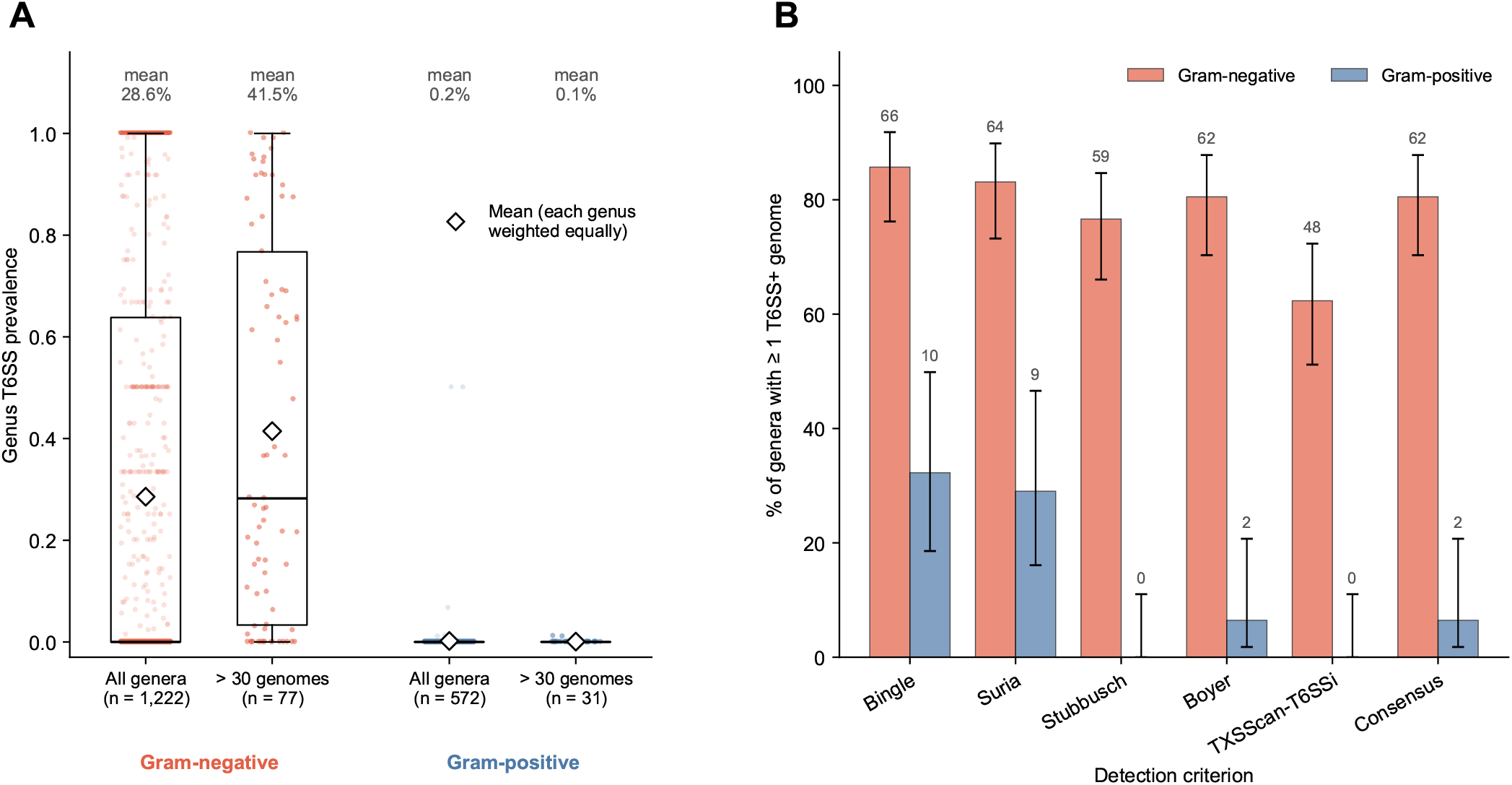
T6SS prevalence by Gram status. (A) Genus T6SS prevalence by the consensus call for all genera with a Gram stain in BacDive (1,222 Gram-negative and 572 Gram-positive genera) and for the well-sampled genera among them, those with more than 30 sampled genomes (77 and 31). Points are genera, boxes show the median and interquartile range (whiskers to 1.5 times the interquartile range), and diamonds and the values above mark the mean, with each genus weighted equally. Gram-negative prevalence is higher among well-sampled genera (41.5% against 28.6%), while Gram-positive prevalence stays near zero (0.1% and 0.2%). (B) Percentage of well-sampled genera with at least one T6SS-positive genome under each criterion and the consensus, with 95% Wilson score intervals; numbers above the bars give the count of genera.

**Fig S6.**
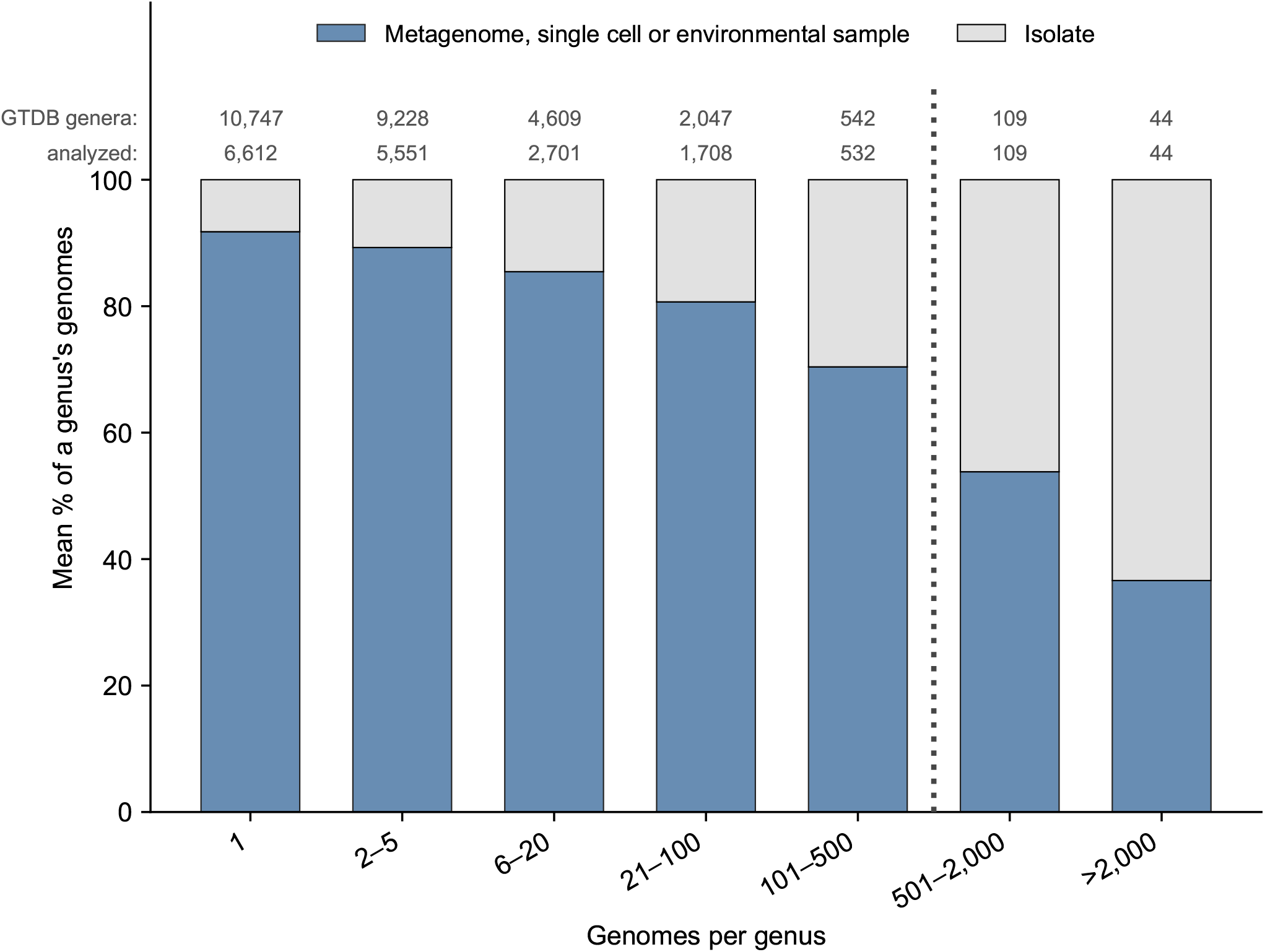
Genus size and the 500-genome sampling threshold. Mean percentage of a genus’s genomes derived from metagenomes, single cells or environmental samples (NCBI genome category) and from isolates, by genus size, for the 27,326 GTDB r226 genera, with each genus weighted equally. Numbers above the bars give the GTDB genera in each bin and, below them, the genera analyzed (17,257 in total: those with at least one sampled genome with protein annotations). The dotted line marks the 500-genome threshold: the 27,173 genera with at most 500 genomes were sampled at 10% (at least one genome), and the 153 genera with more than 500 genomes, which hold 60.4% of all genomes, were sampled at 100 genomes each.

**Fig S7.**
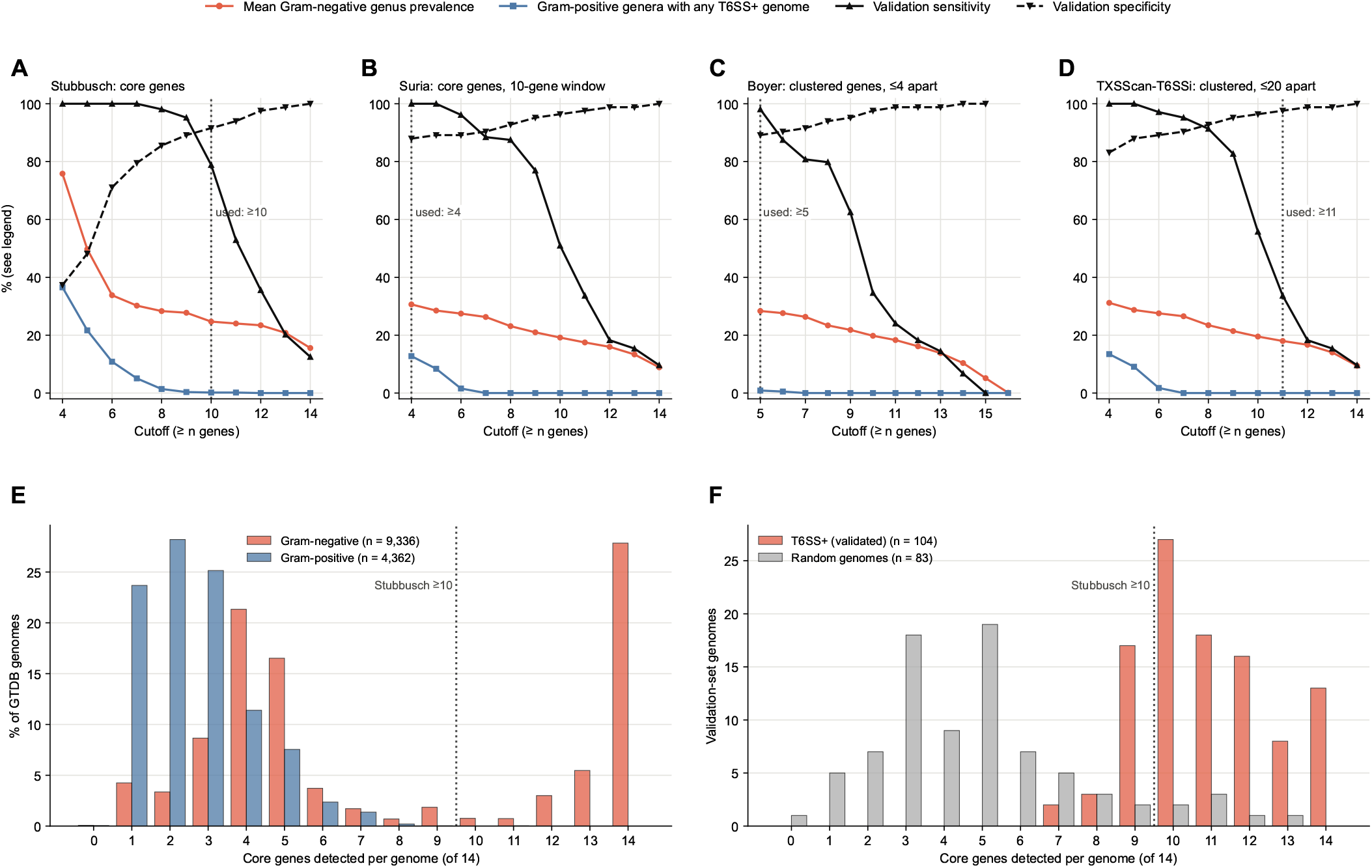
Dependence of prevalence and validation performance on criterion cutoffs. Each count-based criterion calls a genome T6SS-positive when a gene count reaches a cutoff; here every genome is re-called at each cutoff. (A–D) For the Stubbusch (core genes in the genome), Suria (core genes within a 10-gene window), Boyer (T6SS genes in a cluster no more than four genes apart) and TXSScan-T6SSi (core genes in a cluster no more than twenty genes apart) criteria, four quantities as the cutoff changes: mean T6SS prevalence across Gram-negative genera (red); the percentage of Gram-positive genera with at least one T6SS-positive genome (blue), which serves as a false-positive control because Gram-positive bacteria are not expected to encode a T6SS; and sensitivity (solid black) and specificity (dashed black) on the 187-genome validation set (Fig. S3). Dotted lines mark the cutoffs used. Low cutoffs inflate prevalence and call Gram-positive genera T6SS-positive; beyond about seven to eight genes the Gram-negative prevalence changes little and Gram-positive calls fall to near zero, while sensitivity drops steeply above nine to ten genes. Cutoffs start at four core genes (five for Boyer), below which gene positions were not retrieved. (E) Core genes detected per genome (of fourteen), as a percentage of the genomes in each group, for the 13,698 sampled genomes whose genus has a Gram assignment (9,336 from Gram-negative genera and 4,362 from Gram-positive genera). Gram-negative genomes mostly carry either few core genes or nearly all fourteen, with few in between, so cutoffs within this gap give similar calls. Most Gram-positive genomes carry three or fewer, and 23% carry four or more. (F) The same for the validation set: the 104 genomes with experimentally validated T6SSs carry seven to fourteen core genes, and most of the 83 randomly chosen genomes carry fewer than seven. Dotted lines in (E) and (F) mark the Stubbusch cutoff (≥10).

## >Bibliography

1. Stefan Pukatzki, Amy T. Ma, Derek Sturtevant, Bryan Krastins, David Sarracino, William C. Nelson, John F. Heidelberg, and John J. Mekalanos. Identification of a conserved bacterial protein secretion system in Vibrio cholerae using the Dictyostelium host model system. Proceedings of the National Academy of Sciences, 103(5):1528–1533, January 2006. doi: 10.1073/pnas.0510322103. Publisher: Proceedings of the National Academy of Sciences.

2. Joseph D. Mougous, Marianne E. Cuff, Stefan Raunser, Aimee Shen, Min Zhou, Casey A. Gifford, Andrew L. Goodman, Grazyna Joachimiak, Claudia L. Ordoñez, Stephen Lory, Thomas Walz, Andrzej Joachimiak, and John J. Mekalanos. A Virulence Locus of Pseudomonas aeruginosa Encodes a Protein Secretion Apparatus. Science, 312(5779):1526–1530, June 2006. doi: 10.1126/science.1128393.

3. Julia Monjarás Feria and Miguel A. Valvano. An Overview of Anti-Eukaryotic T6SS Effectors. Frontiers in Cellular and Infection Microbiology, 10, October 2020. ISSN 2235-2988. doi: 10.3389/fcimb.2020.584751. Publisher: Frontiers.

4. Benoit Chassaing and Eric Cascales. Antibacterial Weapons: Targeted Destruction in the Microbiota. Trends in Microbiology, 26(4):329–338, April 2018. ISSN 0966-842X, 1878-4380. doi: 10.1016/j.tim.2018.01.006.

5. Lauren Speare, Andrew G. Cecere, Kirsten R. Guckes, Stephanie Smith, Michael S. Wollenberg, Mark J. Mandel, Tim Miyashiro, and Alecia N. Septer. Bacterial symbionts use a type VI secretion system to eliminate competitors in their natural host. Proceedings of the National Academy of Sciences of the United States of America, 115(36):E8528–E8537, September 2018. ISSN 0027-8424. doi: 10.1073/pnas.1808302115.

6. Ramses Gallegos-Monterrosa and Sarah J Coulthurst. The ecological impact of a bacterial weapon: microbial interactions and the Type VI secretion system. FEMS Microbiology Reviews, 45(6):fuab033, June 2021. ISSN 0168-6445. doi: 10.1093/femsre/fuab033.

7. Perla Jazmin Gonzalez Moreno and Michele K. Nishiguchi. The Competitive Edge: T6SS-Mediated Interference Competition by Vibrionaceae Across Marine Ecological Niches. Microorganisms, 13(6):1370, June 2025. ISSN 2076-2607. doi: 10.3390/microorganisms13061370.

8. Jan C. Wohlfarth, Danny Ward, Joana Pereira, and Marek Basler. The type VI secretion system and associated effector proteins. Nature Reviews Microbiology, pages 1–16, November 2025. ISSN 1740-1534. doi: 10.1038/s41579-025-01256-w.

9. Lihua Chen, Yaru Zou, Pengfei She, and Yong Wu. Composition, function, and regulation of T6SS in Pseudomonas aeruginosa. Microbiological Research, 172:19–25, March 2015. ISSN 0944-5013. doi: 10.1016/j.micres.2015.01.004.

10. Ziduo Chen, Yikai Mao, Yinzhou Song, Mengxuan Dou, Ke Shang, Zuhua Yu, Ke Ding, and Songbiao Chen. Refined egoist: The toxin–antitoxin immune system of T6SS. Microbial Pathogenesis, 196:106991, November 2024. ISSN 0882-4010. doi: 10.1016/j.micpath.2024.106991.

11. Elisa T. Granato, William P. J. Smith, and Kevin R. Foster. Collective protection against the type VI secretion system in bacteria. The ISME Journal, 17(7):1052–1062, July 2023. ISSN 1751-7370. doi: 10.1038/s41396-023-01401-4. Publisher: Nature Publishing Group.

12. Tobias Zachs, Jessie James L. Malit, Jingwei Xu, Alexandra Schürch, Shamphavi Sivabalasarma, Phillip Nußbaum, Sonja-Verena Albers, and Martin Pilhofer. ]Archaeal type six secretion system mediates contact-dependent antagonism. Science Advances, 10(46): eadp7088, November 2024. doi: 10.1126/sciadv.adp7088. Publisher: American Association for the Advancement of Science.

13. Patricia Bernal, María A. Llamas, and Alain Filloux. Type VI secretion systems in plant-associated bacteria. Environmental Microbiology, 20(1):1–15, January 2018. ISSN 1462-2912. doi: 10.1111/1462-2920.13956.

14. Astrid K. M. Stubbusch, François J. Peaudecerf, Kang Soo Lee, Lucas Paoli, Julia Schwartzman, Roman Stocker, Marek Basler, Olga T. Schubert, Martin Ackermann, Cara Magnabosco, and Glen G. D’Souza. Antagonism as a foraging strategy in microbial communities. Science, 388(6752):1214–1217, June 2025. doi: 10.1126/science.adr8286. Publisher: American Association for the Advancement of Science.

15. Meiru Si, Chao Zhao, Brianne Burkinshaw, Bing Zhang, Dawei Wei, Yao Wang, Tao G. Dong, and Xihui Shen. Manganese scavenging and oxidative stress response mediated by type VI secretion system in Burkholderia thailandensis. Proceedings of the National Academy of Sciences, 114(11):E2233–E2242, March 2017. doi: 10.1073/pnas.1614902114.

16. Sandra Schwarz, T. Eoin West, Frédéric Boyer, Wen-Chi Chiang, Mike A. Carl, Rachel D. Hood, Laurence Rohmer, Tim Tolker-Nielsen, Shawn J. Skerrett, and Joseph D. Mougous. Burkholderia Type VI Secretion Systems Have Distinct Roles in Eukaryotic and Bacterial Cell Interactions. PLOS Pathogens, 6(8):e1001068, August 2010. ISSN 1553-7374. doi: 10.1371/journal.ppat.1001068.

17. Erick V. S. Motta, Patrick J. Lariviere, Korin R. Jones, Yulin Song, and Nancy A. Moran. Type VI secretion systems promote intraspecific competition and host interactions in a bee gut symbiont. Proceedings of the National Academy of Sciences, 121(44):e2414882121, October 2024. doi: 10.1073/pnas.2414882121.

18. Michael L Weltzer, Pravas C Roy, Jack Govaerts, and Daniel Wall. Aggregative chimeric multicellularity in the absence of lethal kin discrimination. The ISME Journal, 20(1):wrag136, June 2026. ISSN 1751-7362. doi: 10.1093/ismejo/wrag136.

19. Simon B. Otto, Richard Servajean, Alexandre Lemopoulos, Anne-Florence Bitbol, and Melanie Blokesch. Interactions between pili affect the outcome of bacterial competition driven by the type VI secretion system. Current Biology, 34(11):2403–2417.e9, June 2024. ISSN 0960-9822. doi: 10.1016/j.cub.2024.04.041.

20. Jing Jie, Sanwei Gu, Dan Li, Meng Zhang, Zhao-Qing Luo, and Lei Song. The type VI secretion system of Acinetobacter: mechanisms, biology and therapeutic potential. Communications Biology, 9(1):327, February 2026. ISSN 2399-3642. doi: 10.1038/s42003-026-09782-w.

21. Janet Chow and Sarkis K. Mazmanian. A Pathobiont of the Microbiota Balances Host Colonization and Intestinal Inflammation. Cell Host & Microbe, 7(4):265–276, April 2010. ISSN 1931-3128, 1934-6069. doi: 10.1016/j.chom.2010.03.004.

22. Xiaoli Jiang, Hanzeng Li, Jiayue Ma, Hong Li, Xiang Ma, Yanqiong Tang, Juanjuan Li, Xue Chi, Yong Deng, Sheng Zeng, and Zhu Liu. Role of Type VI secretion system in pathogenic remodeling of host gut microbiota during Aeromonas veronii infection. The ISME Journal, 18(1):wrae053, January 2024. ISSN 1751-7362. doi: 10.1093/ismejo/wrae053.

23. Antonio Serapio-Palacios, Sarah E. Woodward, Stefanie L. Vogt, Wanyin Deng, Anna Creus-Cuadros, Kelsey E. Huus, Mihai Cirstea, Madeleine Gerrie, Weronika Barcik, Hongbing Yu, and B. Brett Finlay. Type VI secretion systems of pathogenic and commensal bacteria mediate niche occupancy in the gut. Cell Reports, 39(4), April 2022. ISSN 2211-1247. doi: 10.1016/j.celrep.2022.110731.

24. Jovana Mijatović Scouten Shan-Chi Hsieh, Li-Kang Sung, Yung-Hui Victoria Wen, ChihHorng Kuo, Erh-Min Lai, and Jeff H. Chang. Function, Evolution, and Ecology of Type VI Secretion Systems of Plant-Associated Bacteria. Annual Review of Phytopathology, 63 (Volume 63, 2025):333–356, September 2025. ISSN 0066-4286, 1545-2107. doi: 10.1146/annurev-phyto-121423-084620.

25. Lewis EH Bingle, Christopher M Bailey, and Mark J Pallen. Type VI secretion: a beginner’s guide. Current Opinion in Microbiology, 11(1):3–8, February 2008. ISSN 1369-5274. doi: 10.1016/j.mib.2008.01.006.

26. Giuseppina Mariano, Katharina Trunk, David J. Williams, Laura Monlezun, Henrik Strahl, Samantha J. Pitt, and Sarah J. Coulthurst. A family of Type VI secretion system effector proteins that form ion-selective pores. Nature Communications, 10(1):5484, December 2019. ISSN 2041-1723. doi: 10.1038/s41467-019-13439-0.

27. David Williams. djw533/hamburger, February 2026. original-date: 2019-05-03T10:13:42Z.

28. Young-Jun Park, Kaitlyn D Lacourse, Christian Cambillau, Frank DiMaio, Joseph D Mougous, and David Veesler. Structure of the type vi secretion system tssk–tssf–tssg baseplate subcomplex revealed by cryo-electron microscopy. Nature communications, 9 (1):5385, 2018.

29. Sergey Nazarov, Johannes P Schneider, Maximilian Brackmann, Kenneth N Goldie, Henning Stahlberg, and Marek Basler. zCryo-em reconstruction of type vi secretion system baseplate and sheath distal end. The EMBO journal, 37(4):EMBJ201797103, 2018.

30. Aravindan Ilangovan, Sarah Connery, and Gabriel Waksman. Structural biology of the Gram-negative bacterial conjugation systems. Trends in Microbiology, 23(5):301–310, May 2015. ISSN 0966842X. doi: 10.1016/j.tim.2015.02.012.

31. Donovan H Parks, Maria Chuvochina, Christian Rinke, Aaron J Mussig, Pierre-Alain Chaumeil, and Philip Hugenholtz. GTDB: an ongoing census of bacterial and archaeal diversity through a phylogenetically consistent, rank normalized and complete genome-based taxonomy. Nucleic Acids Research, 50(D1):D785–D794, January 2022. ISSN 0305-1048. doi: 10.1093/nar/gkab776.

32. Donovan H Parks, Pierre-Alain Chaumeil, Aaron J Mussig, Christian Rinke, Maria Chuvochina, and Philip Hugenholtz. GTDB release 10: a complete and systematic taxonomy for 715 230 bacterial and 17 245 archaeal genomes. Nucleic Acids Research, 54(D1): D743–D754, January 2026. ISSN 1362-4962. doi: 10.1093/nar/gkaf1040.

33. Aline L. de Oliveira, Nicolle L. Barbieri, Darby M. Newman, Meaghan M. Young, Lisa K. Nolan, and Catherine M. Logue. Characterizing the Type 6 Secretion System (T6SS) and its role in the virulence of avian pathogenic Escherichia coli strain APECO18. PeerJ, 9: e12631, December 2021. ISSN 2167-8359. doi: 10.7717/peerj.12631. Publisher: PeerJ Inc.

34. Sergio Morgado and Ana Carolina Vicente. Diversity and distribution of Type VI Secretion System gene clusters in bacterial plasmids. Scientific Reports, 12(1):8249, May 2022. ISSN 2045-2322. doi: 10.1038/s41598-022-12382-3. Publisher: Nature Publishing Group.

35. Wenjian Liao, Hai-hua Huang, Qi-sen Huang, Du Fang-Ling, Dan Dan Wei, Wan La-Gen, Tian-xin Xiang, Wei Zhang, and Yang Liu. Distribution of type VI secretion system (T6SS) in clinical Klebsiella pneumoniae strains from a Chinese hospital and its potential relationship with virulence and drug resistance. Microbial Pathogenesis, 162:105085, January 2022. ISSN 0882-4010. doi: 10.1016/j.micpath.2021.105085.

36. Fernando Navarro-Garcia, Fernando Ruiz-Perez, Ángel Cataldi, and Mariano Larzábal. Type VI Secretion System in Pathogenic Escherichia coli: Structure, Role in Virulence, and Acquisition. Frontiers in Microbiology, 10, August 2019. ISSN 1664-302X. doi: 10.3389/fmicb.2019.01965. Publisher: Frontiers.

37. Luke P. Allsopp and Patricia Bernal. Killing in the name of: T6SS structure and effector diversity. Microbiology, 169(7):001367, 2023. ISSN 1465-2080. doi: 10.1099/mic.0.001367. Publisher: Microbiology Society,.

38. Frédéric Boyer, Gwennaële Fichant, Jérémie Berthod, Yves Vandenbrouck, and Ina Attree. Dissecting the bacterial type VI secretion system by a genome wide in silico analysis: what can be learned from available microbial genomic resources? BMC Genomics, 10(1):104, March 2009. ISSN 1471-2164. doi: 10.1186/1471-2164-10-104.

39. Peilin Liu, Awen Yang, Bin Tang, Zhiqian Wang, Zijuan Jian, Yanjun Liu, Jiahui Wang, Baiyun Zhong, Qun Yan, and Wenen Liu. Molecular epidemiology and clinical characteristics of the type VI secretion system in Klebsiella pneumoniae causing abscesses. Frontiers in Microbiology, 14, May 2023. ISSN 1664-302X. doi: 10.3389/fmicb.2023.1181701. Publisher: Frontiers.

40. David J. Williams, Alexandra Hawkins, Ruth E. Hernandez, Giuseppina Mariano, Katharine Mathers, Grant Buchanan, Barnaby J. Stonier, Teresa Inkster, Alistair Leanord, James D. Chalmers, Nicholas R. Thomson, Matthew T. G. Holden, and Sarah J. Coulthurst. Competitive behaviors in Serratia marcescens are coordinately regulated by a lifestyle switch frequently inactivated in the clinical environment. Cell Host & Microbe, 33(2):252–266.e5, February 2025. ISSN 1931-3128. doi: 10.1016/j.chom.2025.01.001.

41. Tingting Dong, Wenjing Yi, Meng Zhang, Ning Zhu, Jing Jie, Zhihan Peng, Lili Jiang, Chunyan Wang, Lei Song, Shucheng Hua, and Qingtian Guan. Comprehensive analysis of the type VI secretion system in Neisseria: identification, distribution, and evolutionary insights. BMC Genomics, 26(1):439, May 2025. ISSN 1471-2164. doi: 10.1186/s12864-025-11615-9.

42. Andrea M. Suria, Stephanie Smith, Lauren Speare, Yuzhou Chen, Iris Chien, Emily Grace Clark, Madelyn Krueger, Alexander M. Warwick, Hannah Wilkins, and Alecia N. Septer. Prevalence and diversity of type VI secretion systems in a model beneficial symbiosis. Frontiers in Microbiology, 13, September 2022. ISSN 1664-302X. doi: 10.3389/fmicb.2022.988044. Publisher: Frontiers.

43. Bertrand Néron, Rémi Denise, Charles Coluzzi, Marie Touchon, Eduardo P. C. Rocha, and Sophie S. Abby. MacSyFinder v2: Improved modelling and search engine to identify molecular systems in genomes. Peer Community Journal, 3, March 2023. ISSN 2804-3871. doi: 10.24072/pcjournal.250.

44. Jianfeng Zhang, Jiahao Guan, Meng Wang, Gang Li, Marko Djordjevic, Cui Tai, Hui Wang, Zixin Deng, Zhaoyan Chen, and Hong-Yu Ou. Secret6 update: a comprehensive resource of bacterial type vi secretion systems. Science China Life Sciences, 66(3):626–634, 2023.

45. Isabel Schober, Julia Koblitz, Joaquim Sardà Carbasse Christian Ebeling, Marvin Leon Schmidt, Adam Podstawka, Rohit Gupta, Vinodh Ilangovan, Javad Chamanara, Jörg Overmann, and Lorenz Christian Reimer. BacDive in 2025: the core database for prokaryotic strain data. Nucleic Acids Research, 53(D1):D748–D756, January 2025. ISSN 1362-4962. doi: 10.1093/nar/gkae959.

46. Van Son Nguyen, Silvia Spinelli, Éric Cascales, Alain Roussel, Christian Cambillau, and Philippe Leone. Anchoring the T6SS to the cell wall: Crystal structure of the peptidoglycan binding domain of the TagL accessory protein. PLoS ONE, 16(7):e0254232, July 2021. ISSN 1932-6203. doi: 10.1371/journal.pone.0254232.

47. Dor Salomon, Herman Gonzalez, Barrett L. Updegraff, and Kim Orth. Vibrio parahaemolyticus Type VI Secretion System 1 Is Activated in Marine Conditions to Target Bacteria, and Is Differentially Regulated from System 2. PLOS ONE, 8(4):e61086, April 2013. ISSN 1932-6203. doi: 10.1371/journal.pone.0061086.

48. Matthieu Barret, Frank Egan, Emilie Fargier, John P. Morrissey, and Fergal O’Gara. Genomic analysis of the type VI secretion systems in Pseudomonas spp.: novel clusters and putative effectors uncovered. Microbiology, 157(6):1726–1739, 2011. ISSN 1465-2080. doi: 10.1099/mic.0.048645-0.

49. Eryn E. Bernardy, Maryann A. Turnsek, Sarah K. Wilson, Cheryl L. Tarr, and Brian K. Hammer. Diversity of Clinical and Environmental Isolates of Vibrio cholerae in Natural Transformation and Contact-Dependent Bacterial Killing Indicative of Type VI Secretion System Activity. Applied and Environmental Microbiology, 82(9):2833–2842, May 2016. doi: 10.1128/AEM.00351-16. Publisher: American Society for Microbiology.

50. Prabha Liyanapathiranage, Naama Wagner, Oren Avram, Tal Pupko, and Neha Potnis. Phylogenetic Distribution and Evolution of Type VI Secretion System in the Genus Xanthomonas. Frontiers in Microbiology, 13, April 2022. ISSN 1664-302X. doi: 10.3389/fmicb.2022.840308.

51. Carlos J. Blondel, Juan C. Jiménez, Inés Contreras, and Carlos A. Santiviago. Comparative genomic analysis uncovers 3 novel loci encoding type six secretion systems differentially distributed in Salmonella serotypes. BMC Genomics, 10(1):354, August 2009. ISSN 1471-2164. doi: 10.1186/1471-2164-10-354.

52. Brent S. Weber, Sarah T. Miyata, Jeremy A. Iwashkiw, Brittany L. Mortensen, Eric P. Skaar, Stefan Pukatzki, and Mario F. Feldman. Genomic and Functional Analysis of the Type VI Secretion System in Acinetobacter. PLOS ONE, 8(1):e55142, January 2013. ISSN 1932-6203. doi: 10.1371/journal.pone.0055142.

53. Guillermo D. Repizo, Stéphanie Gagné, Marie-Laure Foucault-Grunenwald, Vitor Borges, Xavier Charpentier, Adriana S. Limansky, João Paulo Gomes, Alejandro M. Viale, and Suzana P. Salcedo. Differential Role of the T6SS in Acinetobacter baumannii Virulence. PLOS ONE, 10(9):e0138265, September 2015. ISSN 1932-6203. doi: 10.1371/journal.pone.0138265.

54. Kvin Lertpiriyapong, Eric R. Gamazon, Yan Feng, Danny S. Park, Jassia Pang, Georgina Botka, Michelle E. Graffam, Zhongming Ge, and James G. Fox. Campylobacter jejuni Type VI Secretion System: Roles in Adaptation to Deoxycholic Acid, Host Cell Adherence, Invasion, and In Vivo Colonization. PLOS ONE, 7(8):e42842, August 2012. ISSN 1932-6203. doi: 10.1371/journal.pone.0042842.

55. M. Ugarte-Ruiz, R. A. Stabler, L. Domínguez, M. C. Porrero, B. W. Wren, N. Dorrell, and O. Gundogdu. Prevalence of Type VI Secretion System in Spanish Campylobacter jejuni Isolates. Zoonoses and Public Health, 62(7):497–500, 2015. ISSN 1863-2378. doi: 10.1111/zph.12176.

56. Divine Yufetar Shyntum, Stephanus Nicolaas Venter, Lucy Novungayo Moleleki, Ian Toth, and Teresa Ann Coutinho. Comparative genomics of type VI secretion systems in strains of Pantoea ananatisfrom different environments. BMC Genomics, 15(1):163, February 2014. ISSN 1471-2164. doi: 10.1186/1471-2164-15-163.

57. Hasan C. Tekedar, Hossam Abdelhamed, Salih Kumru, Jochen Blom, Attila Karsi, and Mark L. Lawrence. Comparative Genomics of Aeromonas hydrophila Secretion Systems and Mutational Analysis of hcp1 and vgrG1 Genes From T6SS. Frontiers in Microbiology, 9:3216, January 2019. ISSN 1664-302X. doi: 10.3389/fmicb.2018.03216.

58. Luke P. Allsopp, Patricia Bernal, Laura M. Nolan, and Alain Filloux. Causalities of war: The connection between type VI secretion system and microbiota. Cellular Microbiology, 22(3): e13153, March 2020. ISSN 1462-5814. doi: 10.1111/cmi.13153.

59. Rahul Unni, Katherine L. Pintor, Andreas Diepold, and DanielYR 2022 Unterweger. Presence and absence of type VI secretion systems in bacteria. Microbiology, 168(4):001151, 2022. ISSN 1465-2080,. doi: 10.1099/mic.0.001151.

60. Lieven Wittebolle, Massimo Marzorati, Lieven Clement, Annalisa Balloi, Daniele Daffonchio, Kim Heylen, Paul De Vos, Willy Verstraete, and Nico Boon. Initial community evenness favours functionality under selective stress. Nature, 458(7238):623–626, April 2009. ISSN 1476-4687. doi: 10.1038/nature07840.

61. Christopher Zhang, Sayantan Datta, William C. Ratcliff, and Brian K. Hammer. Constitutive expression of the Type VI Secretion System carries no measurable fitness cost in Vibrio cholerae. Ecology and Evolution, 14(3):e11081, March 2024. ISSN 2045-7758. doi: 10.1002/ece3.11081.

62. Boris Taillefer, Julien F. Giraud, and Eric Cascales. No fitness cost entailed by type VI secretion system synthesis, assembly, contraction, or disassembly in enteroaggregative Escherichia coli. Journal of Bacteriology, 205(12):e00357–23, November 2023. doi: 10.1128/jb.00357-23.

63. Sophie Robitaille, Emilia L. Simmons, Adrian J. Verster, Emily Anne McClure, Darlene Royce, Evan M. Trus, Kerry Swartz, Daniel Schultz, Carey D. Nadell, and Benjamin D. Ross. Community composition and the environment modulate the population dynamics of type VI secretion in human gut bacteria, February 2023. Pages: 2023.02.20.529031 Section: New Results.

64. Thibault R. Bongiovanni, Casey J. Latario, Youn Le Cras, Evan Trus, Sophie Robitaille, Kerry Swartz, Danica Schmidtke, Maxence Vincent, Artemis Kosta, Jan Orth, Florian Stengel, Riccardo Pellarin, Eduardo P. C. Rocha, Benjamin D. Ross, and Eric Durand. Assembly of a unique membrane complex in type VI secretion systems of Bacteroidota. Nature Communications, 15(1):429, January 2024. ISSN 2041-1723. doi: 10.1038/s41467-023-44426-1.

65. Francis J. Santoriello, Paul C. Kirchberger, Yann Boucher, and Stefan Pukatzki. Pandemic Vibrio cholerae acquired competitive traits from an environmental Vibrio species. Life Science Alliance, 6(2), February 2023. ISSN 2575-1077. doi: 10.26508/lsa.202201437. Publisher: Life Science Alliance Section: Research Articles.

66. Benjamin Buchfink, Klaus Reuter, and Hajk-Georg Drost. Sensitive protein alignments at tree-of-life scale using diamond. Nature methods, 18(4):366–368, 2021.

67. Stijn Van Dongen. Graph clustering via a discrete uncoupling process. SIAM Journal on Matrix Analysis and Applications, 30(1):121–141, 2008.

68. Sylvain Brohee and Jacques Van Helden. Evaluation of clustering algorithms for protein-protein interaction networks. BMC bioinformatics, 7(1):488, 2006.

69. Hayley B Hassler, Brett Probert, Carson Moore, Elizabeth Lawson, Richard W Jackson, Brook T Russell, and Vincent P Richards. Phylogenies of the 16s rrna gene and its hypervariable regions lack concordance with core genome phylogenies. Microbiome, 10(1):104, 2022.

70. Carolina A Martinez-Gutierrez and Frank O Aylward. Phylogenetic signal, congruence, and uncertainty across bacteria and archaea. Molecular Biology and Evolution, 38(12):5514–5527, 2021.

71. Fabian Sievers and Desmond G Higgins. Clustal omega for making accurate alignments of many protein sequences. Protein Science, 27(1):135–145, 2018.

72. Subha Kalyaanamoorthy, Bui Quang Minh, Thomas KF Wong, Arndt Von Haeseler, and Lars S Jermiin. Modelfinder: fast model selection for accurate phylogenetic estimates. Nature methods, 14(6):587–589, 2017.

73. Bui Quang Minh, Minh Anh Thi Nguyen, and Arndt Von Haeseler. Ultrafast approximation for phylogenetic bootstrap. Molecular biology and evolution, 30(5):1188–1195, 2013.

74. Bui Quang Minh, Heiko A Schmidt, Olga Chernomor, Dominik Schrempf, Michael D Woodhams, Arndt Von Haeseler, and Robert Lanfear. Iq-tree 2: new models and efficient methods for phylogenetic inference in the genomic era. Molecular biology and evolution, 37(5): 1530–1534, 2020.

75. Guangchuang Yu. Using ggtree to visualize data on tree-like structures. Current protocols in bioinformatics, 69(1):e96, 2020.

76. R Core Team et al. R: A language and environment for statistical computing. r foundation for statistical computing, vienna, austria. http://www.R-project.org/, 2016.

77. Hadley Wickham. ggplot2. Wiley interdisciplinary reviews: computational statistics, 3(2): 180–185, 2011.

